# Chronic cigarette smoke exposure masks pathological features of *Helicobacter pylori* infection while promoting tumor initiation

**DOI:** 10.1101/2024.08.05.604297

**Authors:** Maeve T. Morris, M Blanca Piazuelo, I. Mark Olfert, Xiaojiang Xu, Salik Hussain, Richard M. Peek, Jonathan T. Busada

**Author notes:** **Corresponding authors:** Jonathan T. Busada, WVU School of Medicine, Microbiology, Immunology and Cell Biology, 64 Medical Center Drive, P.O. Box 9177, Morgantown, WV 26506.

## Abstract

Gastric cancer is the fifth most common cancer and the fifth leading cause of cancer deaths worldwide. Chronic infection by the bacterium *Helicobacter pylori* is the most prominent gastric cancer risk factor, but only 1-3% of infected individuals will develop gastric cancer. Cigarette smoking is another independent gastric cancer risk factor, and *H. pylori-*infected smokers are at a 2-11-fold increased risk of gastric cancer development, but the direct impacts of cigarette smoke on *H. pylori* pathogenesis remain unknown. In this study, male C57BL/6 mice were infected with *H. pylori* and began smoking within one week of infection. The mice were exposed to cigarette smoke (CS) five days/week for 8 weeks. CS exposure had no notable impact on gross gastric morphology or inflammatory status compared to filtered-air (FA) exposed controls in mock-infected mice. However, CS exposure significantly blunted *H. pylori-*induced gastric inflammatory responses, reducing gastric atrophy and pyloric metaplasia development. Despite blunting these classic pathological features of *H. pylori* infection, CS exposures increased DNA damage within the gastric epithelial cells and accelerated *H. pylori-*induced dysplasia onset in the INS-GAS gastric cancer model. These data suggest that cigarette smoking may clinically silence classic clinical symptoms of *H. pylori* infection but enhance the accumulation of mutations and accelerate gastric cancer initiation.

## Introduction

Gastric cancer remains one of the most common malignancies worldwide and is the 5^th^ leading cause of cancer deaths ^1^. *Helicobacter pylori* infection is the primary risk factor for gastric cancer, contributing to 75% of the global gastric cancer burden ^2^. *H. pylori* infection is extremely prevalent, infecting approximately 50% of the world’s population. However, only 1-3% of infected individuals will develop cancer ^3^. Early detection and diagnosis of gastric cancer is one of the most significant blockades to effective treatment. When diagnosed early, survival rates are relatively high at 73%, but when diagnosed late, survival is abysmal at 6% ^4^. Identifying individuals that are at a high risk of cancer among the more than four billion infected individuals remains a persistent challenge. Defining how environmental risk factors modify *H. pylori* pathogenesis and cancer development has the potential to prioritized patient populations for monitoring and treatments.

Cigarette smoking has long been recognized as a leading risk factor for gastric cancer, increasing the risk of both cardia and distal gastric cancers ^5, 6^. Cigarette smoking intensity and duration directly correlate to increased cancer risk. A recent meta-analysis of over 200 publications found that gastric cancer risk was significantly increased over never-smokers with as few as 5 cigarettes/day and increased non-linearly up to 20 cigarettes/day ^7^. Similarly, gastric cancer risk increased significantly with smoking duration. Cigarette smoke (CS) is a complex mixture of over 4500 known chemicals, including tar, ammonia, carbon monoxide, formaldehyde, acetone, nitrogen oxides, and heavy metals, and includes more than 60 known or potential carcinogens ^8, 9^. Pulmonary studies demonstrate that CS exposure damages the airway, induces DNA damage, and causes immune dysfunction ^10^. Similar effects may occur within the stomach, as smokers have a significantly higher incidence of peptic ulcer disease, intestinal metaplasia, and dysplasia than nonsmokers ^11, 12^. Moreover, *H. pylori-*infected smokers exhibit lower levels of leukocyte infiltration and lower anti-*H. pylori* antibody titers, suggesting that CS impairs gastric inflammatory responses ^13^.

*Helicobacter pylori* infection and cigarette smoking are leading independent gastric cancer risk factors ^14^, and mounting evidence suggests that they positively interact to further increase gastric cancer risk ^14, 15^. *H. pylori-*infected smokers are at 2-11 fold higher risk of gastric cancer than either *H. pylori-*infected nonsmokers or uninfected smokers ^15^. However, despite the increased risk of gastric cancer in *H. pylori-*infected smokers, how CS affects *H. pylori* pathogenesis and damage responses within the gastric epithelium remain unknown. In this study, we investigated how chronic CS exposures impact *H. pylori-*induced gastric inflammation, atrophy, and metaplasia development. Our data indicate that CS impairs gastric inflammatory responses to *H. pylori* infection, limiting overt gastric atrophy and metaplasia development but enhancing DNA damage, suggesting that smokers should receive additional monitoring for *H. pylori* infection and be prioritized for eradication therapy.

## Material and Methods

### Animal Care and Treatment

All mouse studies were performed with approval by the Animal Care and Use Committee at West Virginia University. C57BL/6J mice were purchased from the Jackson Laboratories. INS-GAS mice were provided from Dr. Jim Fox (Massachusetts Institute of Technology). All mice were administered standard chow and water *ab libitum* and maintained in a temperature- and humidity-controlled room with standard 12-hour light/dark cycles.

### H. pylori preparation, mouse infections, and infection confirmation

The *Helicobacter pylori* PMSS1 strain was grown on tryptic soy agar plates (BD Biosciences) with 5% defibrinated sheep blood (Hemostat Labs) and 10 μg/mL vancomycin (Alfa Aesar) under microaerophilic conditions at 37°C. The bacteria were then harvested and transferred to Brucella broth (Research Products International) containing 5% fetal bovine serum (R&D systems) and 10 μg/mL vancomycin and grown overnight at 37°C under microaerophilic conditions with agitation. Bacteria were centrifuged and resuspended in fresh Brucella broth without antibiotics before spectrophotometry and mouse infection. Bacterial numbers were determined by a standard curve of bacterial counts and optical density. Gram staining was used to confirm culture purity. Mice were colonized by oral gavage, receiving two consecutive doses of 10^9^ bacteria spaced 24 hours apart. Mice were colonized 2 days prior to initiating CS exposure. To confirm colonization, biopsies from the gastric corpus and pylorus were homogenized and plated on tryptic soy agar plates with 5% defibrinated sheep blood containing amphotericin, bacitracin, nalidixic acid, and vancomycin. *H. pylori* growth was confirmed by gram staining.

### Cigarette smoke (CS) exposures

Mock and infected mice were exposed to smoke from Marlboro Red cigarettes in whole-body inhalation chambers using the Gram’s Universal smoke machine (Gram Research). Puff volume was 87ml, with a 5-second inhale and 3-second exhale. Mainstream smoke was pumped into the chamber while sidestream fresh air was constantly provided. Chamber total flow rate was 4.3 L/min. The mice were exposed for approximately 2 hours/day, with each cigarette burning for approximately 7 minutes with a 5-minute break between cigarettes; the mice were exposed to 10 cigarettes/day. Carbon monoxide (CO) levels were monitored throughout the smoke exposures using a URAS26 5-Gas Analyzer. CO levels peaked at 145 ppm and returned to an average of 33 ppm during the 5-minute break. Mock-infected and *H. pylori-*infected control mice were exposed to filtered air in parallel with the smoke-exposed groups. Mice were exposed 5 days/week for 2 months.

### ELISA Assay

Serum was collected by cardiac stick 24 hours after the final smoke exposure. Serum cotinine levels were measured by ELISA assay following the manufacturer’s protocol (Calbiotech).

### Tissue preparation

Mice were euthanized by cervical dislocation without anesthesia. Stomachs were opened along the greater curvature and washed in phosphate buffered saline to remove the gastric contents. Half of the gastric corpus was used for flow cytometry. 4 mm biopsy punches were collected from the gastric corpus for RNA and DNA isolation and immediately snap-frozen in liquid nitrogen. The remainder of the gastric corpus was fixed overnight in 4% paraformaldehyde at 4°C and processed for paraffin embedding or cryosectioning.

### Histology

Standard methods were used when performing immunostaining. 5μm stomach cryosections were incubated with antibodies to H+/K+ ATPase (clone 1H9, MBL Life Sciences), MIST1 (Cell Signaling Technologies), CD45 (clone 104; Biolegend), or CD44v9 (Cosmo Bio) for 1 hour at room temperature or overnight at 4°C. Sections were then incubated in secondary antibodies for 1 hour at room temperature. Where indicated, fluorescence conjugated *Griffonia simplicifolia* lectin (GSII; ThermoFisher) was added with the secondary antibodies. Sections were mounted with Vectastain mounting media containing 4’,6-diamidino-2-phenylindole (Vector Laboratories). Images were obtained using a Zeiss 710 confocal laser-scanning microscope (Carl-Zeiss) and running Zen Black (Carl-Zeiss) imaging software.

### RNA and DNA isolation and qRT-PCR

RNA was extracted in TRIzol (ThermoFisher) and precipitated from the aqueous phase using 1.5 volumes of 100% ethanol. The mixture was transferred to an RNA isolation column (Omega Bio-Tek), and the remaining steps were followed according to the manufacturer’s recommendations. Reverse transcription followed by qPCR was performed in the same reaction using the Universal Probes One-Step PCR kit (Bio-Rad Laboratories) and the TaqMan primers: *Ppib* (Mm00478295_m1)*, Aqp5* (Mm00437578_m1), and *Gkn3* (Mm01183934_m1) (all from Thermofisher). For DNA isolation, gastric tissue was disrupted in 0.5M NaOH at 98°C, then diluted to a final concentration of 100mM Tris-HCl. DNA was purified using a DNA Clean & Concentrator kit (Zymo Research). qPCR was performed using the TaqMan Fast Advanced Master Mix (Thermofisher) and the Taqman primers: *H. pylori* 23s (forward: AACAAGTACCGTGAGGGAAAG; reverse: GCAGTCCATCACCCTGATAAA; probe: /56FAM/AACCGCAGT/ZEN/GAGCGGAGTGAAATA/3IABkFQ/) and *Ppib* (Mm00291049_cn; Thermofisher). *Hp23s* cycle threshold values were normalized to mouse *Ppib* DNA. All qPCR reactions were performed on the QuantStudio 6 Real-Time PCR system (ThermoFisher).

### RNAseq analysis

RNA was isolated from the gastric corpus of *H. pylori-infected* mice after 2 months of exposure to FA or CS. Admera Health performed libraries and sequencing. Libraries were prepared using a NEB ultraII directional kit (New England Biolabs), and 40 million paired-end reads were collected per library using an Illumina HiSeq (Illumina). Reads were aligned to the UCSC mm10 reference genome using STAR with default parameters. The quantification results from featureCounts were then analyzed with the Bioconductor package DESeq2. GSEA analysis was performed using GSEA v4.3.2 software. Genes were preranked based on their log fold change. The significance of the enrichment score was assessed using 1000 permutations. The resulting enriched pathways were visualized using the Cytoscape (v3.10.2) Enrichment Map plugin using a q>0.05 cutoff.

### Flow Cytometry

Harvested tissue was diced and washed in Hanks Balanced Salt Solution without Ca^2+^ or Mg^2+^ containing 5 mM HEPES, 5 mM EDTA, and 5% FBS at 37°C for 20 minutes. Tissue pieces were then washed briefly in Hanks Balanced Salt Solution with Ca^2+^ or Mg^2+^ and then digested in 1mg/ml collagenase (Worthington Biochemical Corporation) for 30 minutes at 37°C. After digestion, tissue fragments were pushed through a 100μM strainer and then rinsed through a 40μM strainer before debris was removed through an Optiprep (Serumwerk) density gradient. Fc receptors were blocked with TruStain (Biolegend), then stained with primary antibodies: CD45.2 (clone 104), CD3e (clone 145-2C11), B220 (clone RA3-6B2), CD4 (clone GK1.5), CD8a (clone 53-6.7) CD11b (clone M1/70), MCHII (clone M5/114.15.2), Ly6G (clone 1A8), F4/80 (clone BM8) (all from Biolegend), and SIGLECF (clone E50-2440; BD Bioscience) for 20 minutes on ice. Actinomycin D (A1310, Invitrogen) was used to label dead cells. Samples were analyzed on a Cytek Aurora spectral flow cytometer (Cytek Biosciences). Flow cytometry analysis was performed using Cytobank (Beckman Coulter).

### Statistical Analysis

All error bars are ± SD of the mean. The sample size for each experiment is indicated in the figure legends. Experiments were repeated a minimum of two times. A one-sample t-test was used when comparing one group. An unpaired t-test was used when comparing two groups. A one-way analysis of variance with the post hoc Tukey t-test was used when comparing three or more groups. Statistical analysis was performed using GraphPad Prism 10 software. Statistical significance was set at *p*≤0.05. Specific *P* values are listed in the figure legends.

## Results

### Cigarette smoke exposure blunts the gastric inflammatory response to H. pylori

*H. pylori* and CS are leading gastric cancer risk factors, but the impacts of CS on *H. pylori* pathogenesis are unknown. To investigate how chronic CS exposure impacts the stomach and *H. pylori* pathogenesis, male C57BL/6J mice were infected with 10^9^ CFUs of the *H. pylori* cagA+ PMSS1 strain ^16^. Two days after colonization, infected and mock-challenged controls were exposed to smoke from Marlboro Red cigarettes in whole-body inhalation chambers using the Gram Universal Smoke Machine. Mock and *H. pylori-*infected control mice were exposed to filtered air (FA) in parallel with the CS-exposed groups. Mice were exposed to CS 5 days/week for 2 months (Figure 1A). To evaluate CS exposure efficacy, serum cotinine levels were measured by ELISA assay immediately after the final smoke exposure. Cotinine levels were 30±16 ng/mL in CS-exposed mice and were undetectable in FA-exposed (Figure 1B). To investigate the effects of CS exposure on the stomach, mice were euthanized immediately after the final smoke exposure. Stomachs were opened along the greater curvature and washed. Chronic CS exposure caused distinct erythema of the proximal stomach and yellowing of the gastric mucosa (Figure 1C), reflecting irritation of the gastric mucosa and direct exposure to CS residue, respectively. To assess whether CS impacted *H. pylori* colonization, relative bacterial density was assessed by qPCR using *H. pylori-*specific primers. DNA was isolated from gastric corpus biopsies, and the qPCR CT-values were normalized to mouse *Ppib.* CS did not significantly impact bacterial density (Figure 1D).

**Figure 1.**
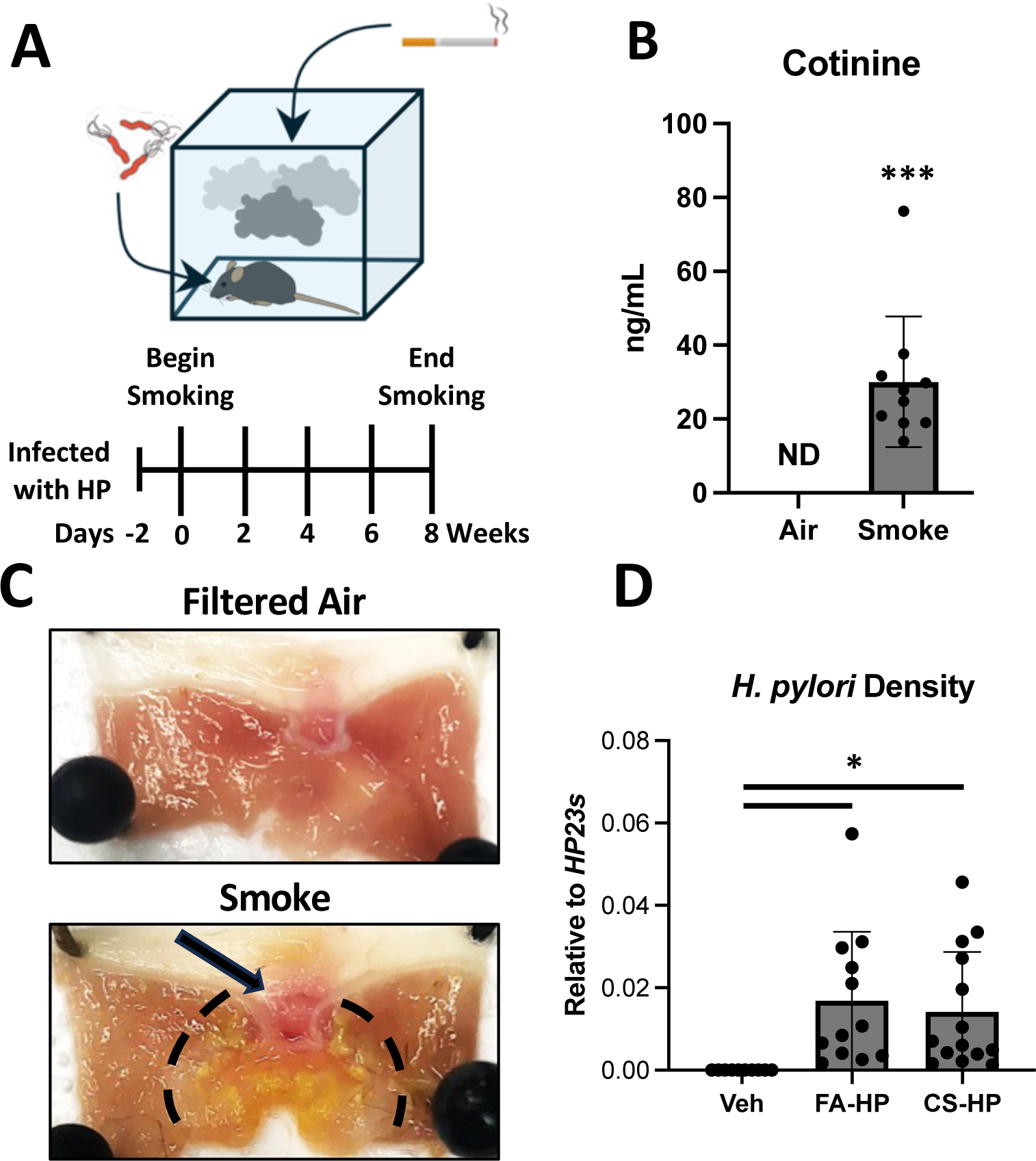
Cigarette smoke residue directly contacts the gastric mucosa. (A) Experimental design. Mice were infected with *H. pylori* and exposed to cigarette smoke 5 days/week for 8 weeks. (B) Serum cotinine levels immediately following final cigarette smoke exposure. (C) Gross gastric morphology of mock-infected mice exposed to either filtered air (FA) or cigarette smoke (CS) for two months. Erythema of the proximal stomach and yellowing of the gastric mucosa are denoted by a hashed line. (D) qRT-PCR for *H. pylori* 23s DNA normalized to mouse *Ppib*. *p≤0.05, **p≤0.01, ***p≤0.001.

Cigarette smoking exerts wide-ranging impacts on immune function ^17^. *H. pylori* colonization is well-known to induce chronic gastric inflammation consisting of prominent T cell, monocytic, and granulocytic infiltrates, which damage the gastric epithelium and drive gastric cancer initiation ^18^. Next, we investigated how CS and *H. pylori* interactions impacted leukocyte recruitment. Immunostaining of mock-infected mice for the common leukocyte antigen CD45 revealed that CS exposure did not induce a noticeable alteration in gross leukocyte recruitment compared to FA-exposed controls (Figure 2A). As expected, *H. pylori* colonization increased CD45+ cells within the gastric corpus. Surprisingly, CS exposure reduced the gross leukocyte recruitment induced by *H. pylori* infection (Figure 2A). Next, we analyzed the gastric corpus immune infiltrate by flow cytometry (Figure 2B). *H. pylori* infection induced significant leukocyte infiltration in FA-exposed versus uninfected controls, but CS exposure significantly reduced *H. pylori-*mediated leukocyte recruitment to the gastric corpus (Figure 2C). Similarly, *H. pylori* significantly induced infiltration of T cells, B cells, Macrophages, Eosinophils, and Neutrophils. CS exposure reduced the infiltration of all immune cell subsets, with the exception of neutrophils (Figure 2C). While total T cell recruitment was reduced by CS exposure, the CD4/CD8 T cell ratios were not changed (Figure 3D). These data demonstrate that CS exposure does not impact *H. pylori* density but significantly impairs the gastric inflammatory response to *H. pylori*.

**Figure 2.**
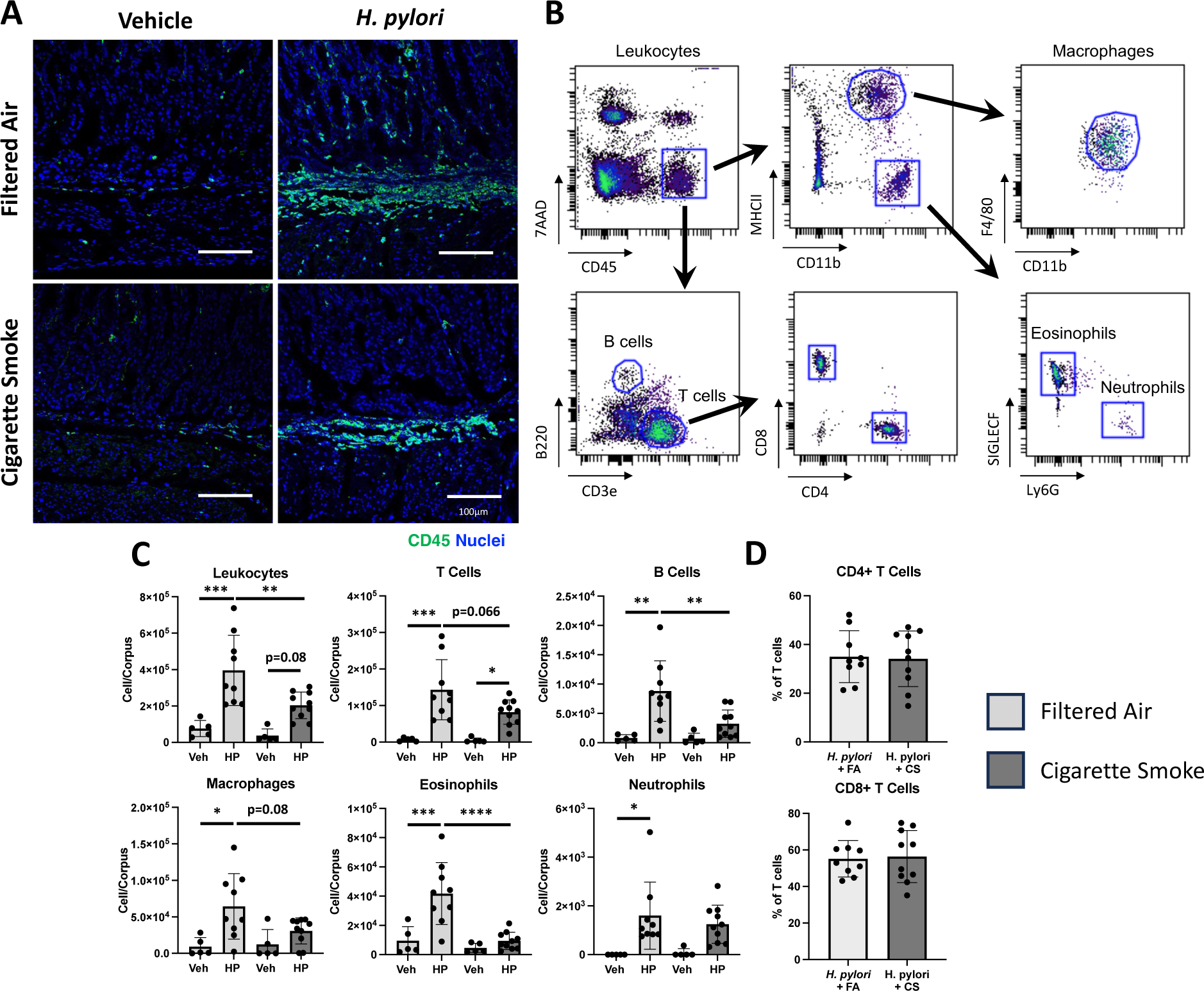
Cigarette smoke limits *H. pylori*-induced gastric inflammation. (A) Representative immunostaining of the common leukocyte antigen CD45 (green). Nuclei are stained in blue. (B) Flow cytometry gating strategy. (C-D) Flow cytometry analysis of cells isolated from the gastric corpus. n≥5 for vehicle groups and n=10 for *H. pylori*-infected groups. Scale bars=100 µm. *p≤0.05, **p≤0.01, ***p≤0.001, and ****p≤0.0001.

**Figure 3.**
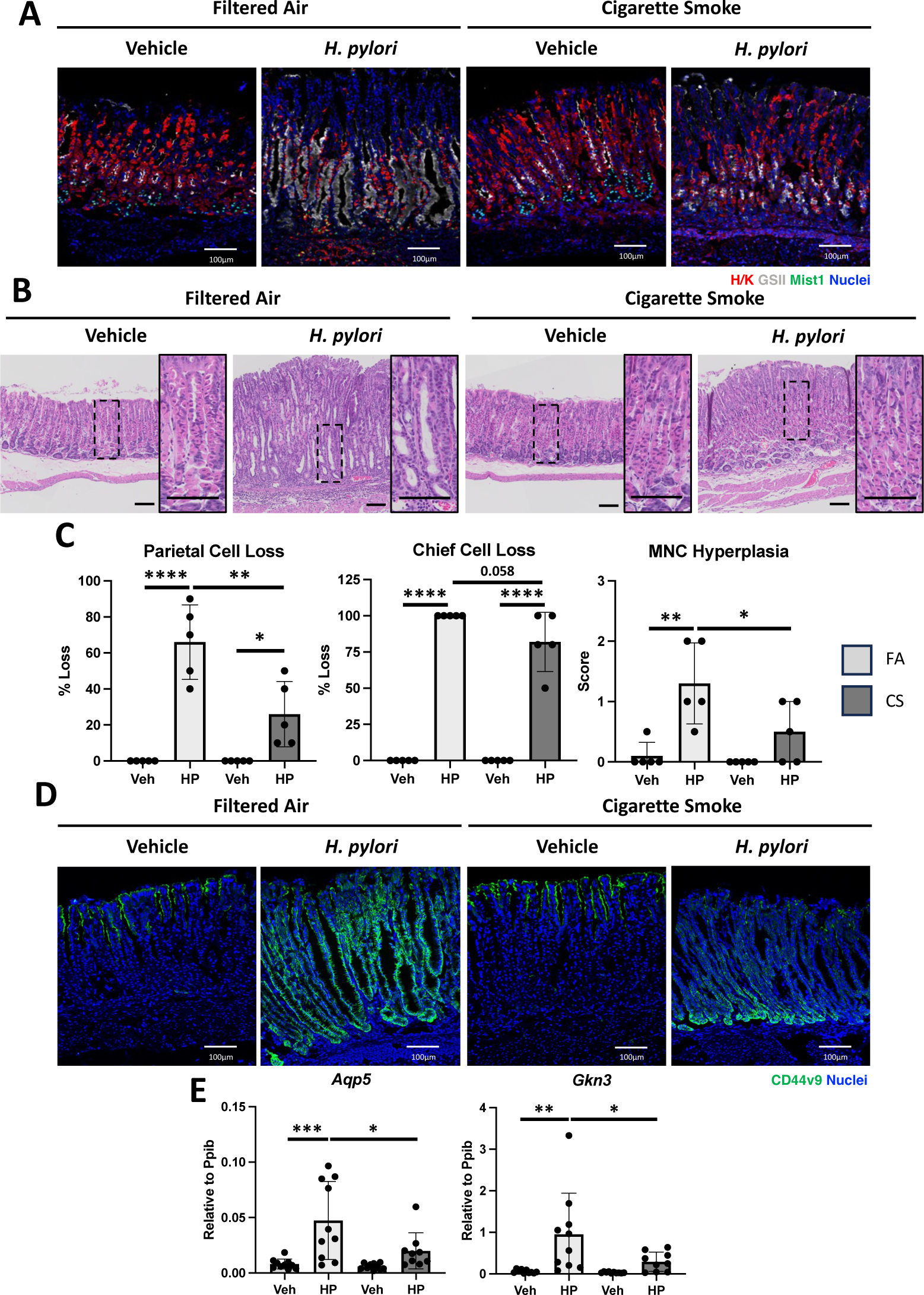
Cigarette smoke exposure dampens *H. pylori*-induced gastric atrophy and metaplasia. (A) Representative immunostaining for parietal cells (red), mucous neck cells (white), and chief cells (green). Nuclei are stained in blue. (B) Representative H&E micrographs of the gastric corpus. (C) Blinded histologic scoring for parietal cell loss, chief cell loss, and mucous neck cell atrophy. (D) Representative immunostaining for the pyloric metaplasia marker CD44 variant 9 (green). Nuclei are stained blue. (E) qRT-PCR for the indicated pyloric metaplasia markers using RNA isolated from the gastric corpus. n≥5. Scale bars=100 µm. *p≤0.05, **p≤0.01, ***p≤0.001, and ****p≤0.0001.

### Cigarette smoke exposure protects from H. pylori-induced gastric atrophy and metaplasia development

Chronic gastric inflammation damages the mucosa, driving atrophy and metaplasia development. Immunostaining for the major corpus lineages parietal cells, chief cells, and mucous neck cells and pathologist scoring of H&E micrographs demonstrated that CS exposure did not impact gastric corpus morphology in mock-infected controls (Figure 3A-C). *H. pylori* infection induced widespread gastric atrophy and significant loss of parietal cells and chief cells and mucous neck cell hyperplasia in FA-exposed controls (Figure 3A-C). In contrast, gastric atrophy and mucous neck cell hyperplasia were significantly attenuated in *H. pylori-*infected mice exposed to CS, consistent with the reduced leukocyte infiltration described above. Gastric atrophy is a precursor of pyloric metaplasia (PM; also called spasmolytic polypeptide-expressing metaplasia), which develops as a healing response to gastric epithelial damage ^19, 20^. Therefore, we assessed pyloric metaplasia development by immunostaining for CD44v9 and performing qRT-PCR for *Aqp5* and *Gkn3*. CD44v9 staining was widespread and intense within the base of the gastric glands (Figure 3D), and *Aqp5* and *Gkn3* transcript levels were significantly elevated in *H. pylori-*infected mice compared to mock-infected controls (Figure 3E). In contrast, CS exposure dramatically reduced CD44v9 expression and significantly reduced *Aqp5* and *Gkn3* expression. These results demonstrate that CS exposure blunts gastritis, gastric atrophy, and metaplasia development, which are classic pathological features associated with *H. pylori* infection.

### Cigarette smoke exposure enhances gastric DNA damage and accelerates dysplasia development

Cigarette smoke contains more than 60 known or suspected carcinogens, and gastric tumors from smokers have increased levels of DNA damage and mutations within the stomach ^8^. Next, we performed bulk RNAseq of the gastric corpus of *H. pylori-*infected mice exposed to FA or CS for two months. Gene Set Enrichment Analysis (GSEA) was used to analyze the list of expressed genes using the REACTOME and BIOCARTA databases. Next, the GSEA results were filtered and functionally clustered by Cytoscape. Gene set clusters for Cell Damage, P53 signaling, DNA repair, Apoptosis, and Cell Death were selected for visualization and were dramatically enriched in the *H. pylori-infected* mice exposed to CS (Figure 4A). These data suggest that oxidative stress and DNA damage are enhanced in stomachs exposed to CS. Therefore, we investigated how chronic CS exposure affected DNA damage within the stomach by immunostaining for the phosphorylated histone variant H2AX as a marker of double-strand DNA breaks. The number of p-H2AX positive cells was equivalent between air-exposed and CS-exposed mice in mock-infected controls (Figure 4B-C). *H. pylori* infection significantly increased p-H2AX positive cells in air-exposed and CS-exposed mice. However, the number of p-H2AX positive cells was significantly higher in CS-exposed mice, indicating that CS synergized with *H. pylori* infection to enhance DNA damage. Finally, we investigated how CS impacts the development of preneoplastic lesions using the INS-GAS mouse model of *H. pylori-*induced gastric cancer. Male INS-GAS mice were colonized with *H. pylori* and exposed to FA or CS for 2 months. Pathologist scoring of H&E micrographs demonstrated a strong trend of increased dysplasia scores in mice exposed to CS (Figure 4D). These data suggest that CS exposure increases gastric DNA damage and accelerates the development of *H. pylori-*mediated gastric preneoplastic lesions.

**Figure 4.**
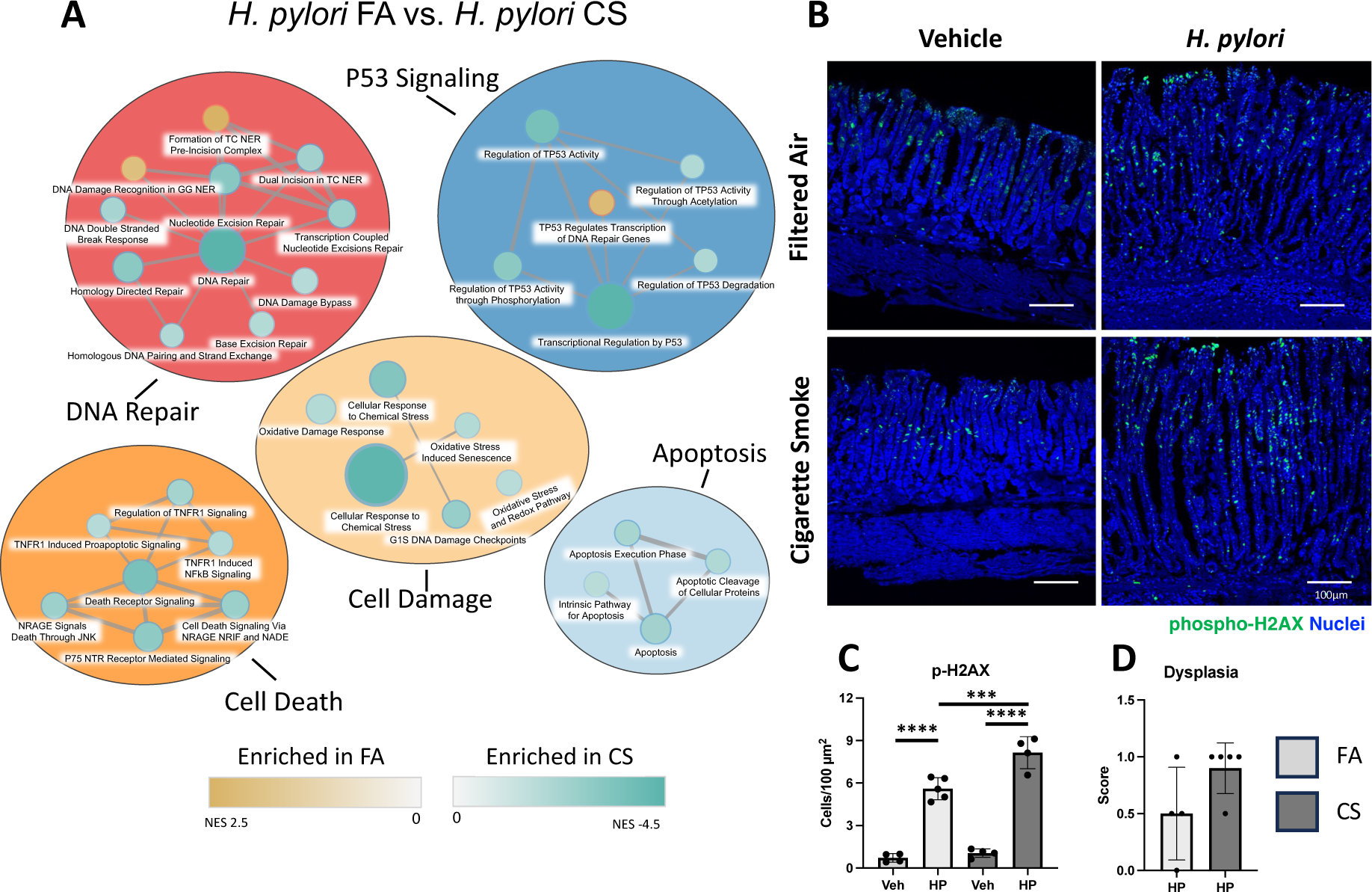
Cigarette smoke exposure increases markers of DNA damage and cancer. (A) RNAseq analysis of the comparing *H. pylori* + FA to *H. pylori +* CS. Gene set enrichment analysis (GSEA) results visualized and clustered by Cytoscape. n=3. (B) Representative immunostaining for phosphorylated gamma histone 2AX (green). Nuclei stained in blue. (C) Quantitation of cells immunostained for phosphorylated gamma histone 2AX. n≥4. (D) Blinded histologic scoring for dysplasia in INS-GAS mice. Scoring scale: 0.5 = metaplasia, 1.0 = dysplasia, 2.0 = carcinoma in situ, 3.0 = invasive adenocarcinoma. N≥4. Scale bars=100 µm. ***p≤0.001, and ****p≤0.0001.

## Discussion

*Helicobacter pylori* infection is the leading gastric cancer risk factor, and bacterial eradication is highly effective at reducing gastric cancer risk ^21^. However, in the United States and other western countries, *H. pylori* screening only occurs in individuals with associated symptoms or those with predisposing gastric cancer risk factors ^22^. Smokers are at increased risk of developing gastric cancer and this risk is compounded by *H. pylori* infection ^15, 23^. However, smokers are not routinely screened for *H. pylori,* nor for preneoplastic lesions in the United States ^24^. Our results demonstrate that CS exposure impairs the inflammatory response to *H. pylori* and blunts the classical pathological features of *H. pylori* infection including gastric atrophy and metaplasia. Simultaneously, CS exposure enhances gastric DNA damage and accelerates dysplasia development. These data indicate that smoking may clinically silence chronic *H. pylori* infections by suppressing atrophic gastritis and metaplasia, but concurrently enhance cancer initiation by increasing mutational burden. Our results suggest that smokers should be screened for *H. pylori* infection, especially if they have additional *H. pylori* risk factors such as low socioeconomic status or are part of ethnic groups with elevated infection rates.

*Helicobacter pylori* infection initiates a histopathological progression called the Correa cascade where inflammation drives gastric atrophy, metaplasia, dysplasia, and eventually results in gastric adenocarcinoma ^25^. Inflammation is critical for driving the development of these pathological features of *H. pylori* infection. Disruption of the T cell compartment effectively eliminates the gastric atrophy and metaplasia associated with *Helicobacter felis* infection in mice ^26^. CD4+ Helper T cells are the primary effector cells of gastric atrophy ^27^. Analysis of bronchiolar lavage fluid and peripheral blood from never smokers, former smokers, and current smokers revealed reduced CD4+ T cells in former and current smokers, while the number of CD8+ T cells were elevated ^28^. We observed that CS blunted total T cell recruitment but that the CD4/CD8 T cell ratio was unchanged. Mice infected with the PMSS1 *H. pylori* strain develop CD8+ biased T cell responses ^29^, which may explain why CS did not further increase CD8 T cell recruitment. Smoke can also induce macrophage dysfunction by reducing phagocytosis and blocking the expression of cytokines and chemokines ^30^. We recently reported that dysfunctional macrophages reduced gastric T cell recruitment during *H. pylori* infection ^31^. Here, we saw that gastric macrophage expansion was blunted by CS-exposure, raising the possibility that impaired macrophages reduced T cell recruitment. Our results agree with gastric biopsies from *H. pylori-* infected smokers who exhibited reduced immune cell infiltrates compared to *H. pylori* infected non-smokers ^13^. We also observed that CS exposure reduced gastric atrophy and pyloric metaplasia development. These *H. pylori-*induced lesions depend on strong anti-*H. pylori* immune responses. Th1-biased responses induce oxyntic atrophy through INFG targeting parietal cells ^32^, while IL13 drive pyloric metaplasia maturation by promoting chief cell transdifferentiation ^33^. CS impairs immunity to bacterial infections in the lungs and reduces IFNG production in mice following oropharyngeal challenge with *Haemophilus influenzae* ^34^. Moreover, CS exposure impairs activation of IL13 gene signatures in pulmonary epithelial cells ^35^. Pyloric metaplasia initially develops in response to gastric glandular damage and likely serves as a healing mechanism ^19, 20^. However, chronic damage and pyloric metaplasia has been observed to transition to intestinal metaplasia and dysplasia in *H. pylori-*infected Mongolian gerbils ^36^. The reduced pyloric metaplasia in CS-exposed mice is likely a consequence of the reduced gastric atrophy and does not indicate a slowing in tumor initiation. Future studies should address the mechanisms of how smoke exposure impairs gastric immunity and epithelial damage responses during *H. pylori* infection.

While gastric atrophy and pyloric metaplasia was reduced by CS, DNA damage and dysplasia in the INS-GAS gastric cancer mice was enhanced. Cigarette smoke is a complex mixture of over 4500 chemicals including more than 60 known carcinogens ^8^. Nicotine and tar compromise immunity to infections and tumor cells while a host of carcinogens cause DNA damage and promote the development of a host of cancer associated with smoking. Gastric tumors from smokers have higher mutational burden that tumors from non-smokers ^37^. The exposure routes taken by these pro-tumorigenic components of CS are poorly understood. CS-associated gasses and particles can be absorbed through the lungs leading to systemic exposure. In addition, ingested particles and CS residue can lead to direct exposure within the gut. Up to 50% of CS particle deposition after a nose-only exposure resides in the digestive tract with the highest concentration in the stomach and esophagus and remains there for several hours ^38^. We found that CS causes macroscopic discoloration of that gastric mucosa and erythema of the proximal stomach adjacent to the esophagus. These findings are consistent with the elevated risk of proximal and cardia tumors in smokers ^39^.

This is the first study to report how CS modifies *H. pylori* pathogenesis. While this work has important implication in screening smokers for *H. pylori,* this study also has limitations. We only utilized a modest CS exposure for a relatively short period. Since gastric cancer risk increases with smoking intensity and duration ^40^, other exposure levels and durations should be considered, especially, long-term studies, to assess tumorigenesis. In addition, we utilized whole-body inhalation exposure to assess the aggregate effects of smoke (i.e., inhalation as well as ingestion). Future studies should also determine the direct and indirect effects of smoking on the stomach and *H. pylori* pathogenesis. Finally, this study utilized male mice. While men are both more likely to smoke than women and are diagnosed with gastric cancer at more than twice the rate of women. Future studies should assess how CS impacts gastric carcinogenesis in both males and females. Overall, this study emphasizes the complex interplay between cigarette smoke, *H. pylori* infection, and gastric carcinogenesis, shedding light on potential mechanisms underlying the elevated risk of gastric cancer in smokers and highlighting the importance of targeted interventions and surveillance in this population.

## Acknowledgments

The authors thank Timothy Nurkiewicz and the West Virginia University Center for Inhalation Toxicology for technical assistance.

## Abbreviations

CS: Cigarette Smoke
FA: Filtered Air
CFUs: Colony Forming Units
PMSS1: Pre-mouse Sydney Strain

## Author Contributions

M.T.M. and J.T.B planned the study. M.T.M. and J.T.B performed experiments. M.B.P performed histological evaluation and XX analyzed the RNAseq data. I.M.O, S.H. and R.M.P. provided essential expertise. M.T.M and J.T.B analyzed all data and drafted the manuscript. All authors reviewed and approved the final manuscript.

